# Early life microbial succession in the gut follows common patterns in humans across the globe

**DOI:** 10.1101/2024.07.25.605223

**Authors:** Guilherme Fahur Bottino, Kevin S. Bonham, Fadheela Patel, Shelley McCann, Michal Zieff, Nathalia Naspolini, Daniel Ho, Theo Portlock, Raphaela Joos, Firas S. Midani, Paulo Schüroff, Anubhav Das, Inoli Shennon, Brooke C. Wilson, Justin M. O’Sullivan, Robert A. Britton, Deirdre M. Murray, Mairead E. Kiely, Carla R. Taddei, Patrícia C. B. Beltrão-Braga, Alline C. Campos, Guilherme V. Polanczyk, Curtis Huttenhower, Kirsten A. Donald, Vanja Klepac-Ceraj

## Abstract

Characterizing the dynamics of microbial community succession in the infant gut microbiome is crucial for understanding child health and development, but no normative model currently exists. Here, we estimate child age using gut microbial taxonomic relative abundances from metagenomes, with high temporal resolution (±3 months) for the first 1.5 years of life. Using 3,154 samples from 1,827 infants across 12 countries, we trained a random forest model, achieving a root mean square error of 2.61 months. We identified key taxonomic predictors of age, including declines in *Bifidobacterium* spp. and increases in *Faecalibacterium prausnitzii* and Lachnospiraceae. Microbial succession patterns are conserved across infants from diverse human populations, suggesting universal developmental trajectories. Functional analysis confirmed trends in key microbial genes involved in feeding transitions and dietary exposures. This model provides a normative benchmark of “microbiome age” for assessing early gut maturation that can be used alongside other measures of child development.

## Introduction

The human gut microbiome is a complex ecosystem consisting of diverse microorganisms that interact with each other and form tight partnerships with their host. These are crucial for several physiological processes, including digestion, metabolism, and immune function^1^. The first major colonization event of an infant’s gastrointestinal tract happens at birth, and microbial succession continues over the first few years of life^2,3^. Age-dependent aspects of this succession are shaped by a combination of natural history and environmental exposures, such as breastfeeding behavior and the introduction of solid food^4,5^. Altered colonization events, especially in early life, may have significant implications on a child’s health, including the development of inflammatory disorders (e.g., allergies and asthma), metabolic disease (e.g., diabetes), neurocognitive outcomes, and other chronic conditions^6,7^.

Specific microbial taxa tend to proliferate at different stages during early infancy^8^. Initial gastrointestinal tract colonizers include microorganisms capable of metabolizing human milk oligosaccharides or scavenging simple molecules^9^. The later introduction of a solid, complex and diverse diet brings opportunity for more fastidious colonizers and a more diverse community^10^. Recurring patterns of colonization and microbial succession across different life stages, from birth to late life and death^11–15^, have shown consistent links between chronology and microbiome development.

These chronology-based approaches have been used to describe the phenotypic implications of an underdeveloped gut microbiome. Studies suggest that when the gut microbial community does not match the expected stage for a child’s age, there can be significant health associations, particularly with growth and immune function^16,17^. This underdevelopment may respond to and contribute to a cycle of poor health and malnutrition, potentially affecting various aspects of the child’s physiology and behavior^18,19^. To measure this temporal mismatch, two things are necessary: a reference developmental trajectory of the gut microbiome in early life and a way to measure a subject’s deviation from such trajectory. One possible solution is to develop age estimation models using gut microbial communities sequenced across large and diverse cohorts. Those models can be trained to accurately produce an estimate of host age that can then be compared with the age at sample collection^17^. Following this approach, links between model outputs and health outcomes in childhood have been reported in multiple areas^20,21^.

Despite showing promise, existing age models face several challenges to be applied in early childhood. Most existing models in this age range utilize data from 16S rRNA gene amplicon sequencing to estimate gut microbiome maturation^17,22^ but this provides only a limited taxonomic resolution as closely related taxa are often binned together^23–25^. Most quantitative age models focus on aging^26–29^ and span large age ranges that either exclude early childhood, or lack the necessary temporal resolution to produce meaningful predictions within the first year of life. Many models that account for early microbiome development with age do not produce a numeric age estimate, instead relying on unsupervised learning and qualitative predictions or associations^30^. Models also tend to be trained on individual cohorts and not validated on external populations, and cross-geographic analyses^31,32^ have been lacking. In recent years, shotgun metagenomic sequencing data has become available from appropriately powered and diverse populations^3^, but these datasets have not yet been incorporated into multi-site age models. Therefore, there is an opportunity and need to develop a comprehensive, global-scale quantitative age model focused on early childhood.

Here, we present such a model for age estimation developed using gut microbial taxonomic relative abundances, with high temporal resolution for the first 1.5 years of life. This model incorporates a large and geographically diverse population, comprising 3,154 shotgun-sequenced samples from 12 countries spanning Africa, Europe, Asia and America.

## Results

### Global metagenomes enable large-scale meta-analysis

We investigated developmental trajectories of the infant gut microbiome using a pooled dataset combining 3,154 stool samples sequenced with shotgun metagenomic sequencing from 1,827 healthy individuals obtained from 12 studies. The metagenomes spanned 12 countries from 4 continents (**Table 1**, **Fig. 1A**). All samples that matched inclusion criteria (see **Methods**) collected between ages 2-18 months (mean = 7.90 mo, SD = 3.99 mo) were incorporated into the model, resulting in a slight overrepresentation of younger samples (ages 2-4 months, **Fig. 1B**, Supplementary Fig. 1). Building the analysis dataset from a wide array of global sources enabled us to include a significant portion of data from low- and middle-income countries (LMICs), representing approximately 46 % of our total sample pool. The 1kD Wellcome LEAP effort contributed a total of 1,817 samples that have not been used previously in age-related studies. 427 of those samples were collected by the Khula study in South Africa^33^ and have not been published before. These 1kD-LEAP samples are slightly younger (mean = 6.86 mo, SD = 3.55 mo), and the majority (80.57 %) are from LMICs.

**Figure 1.**
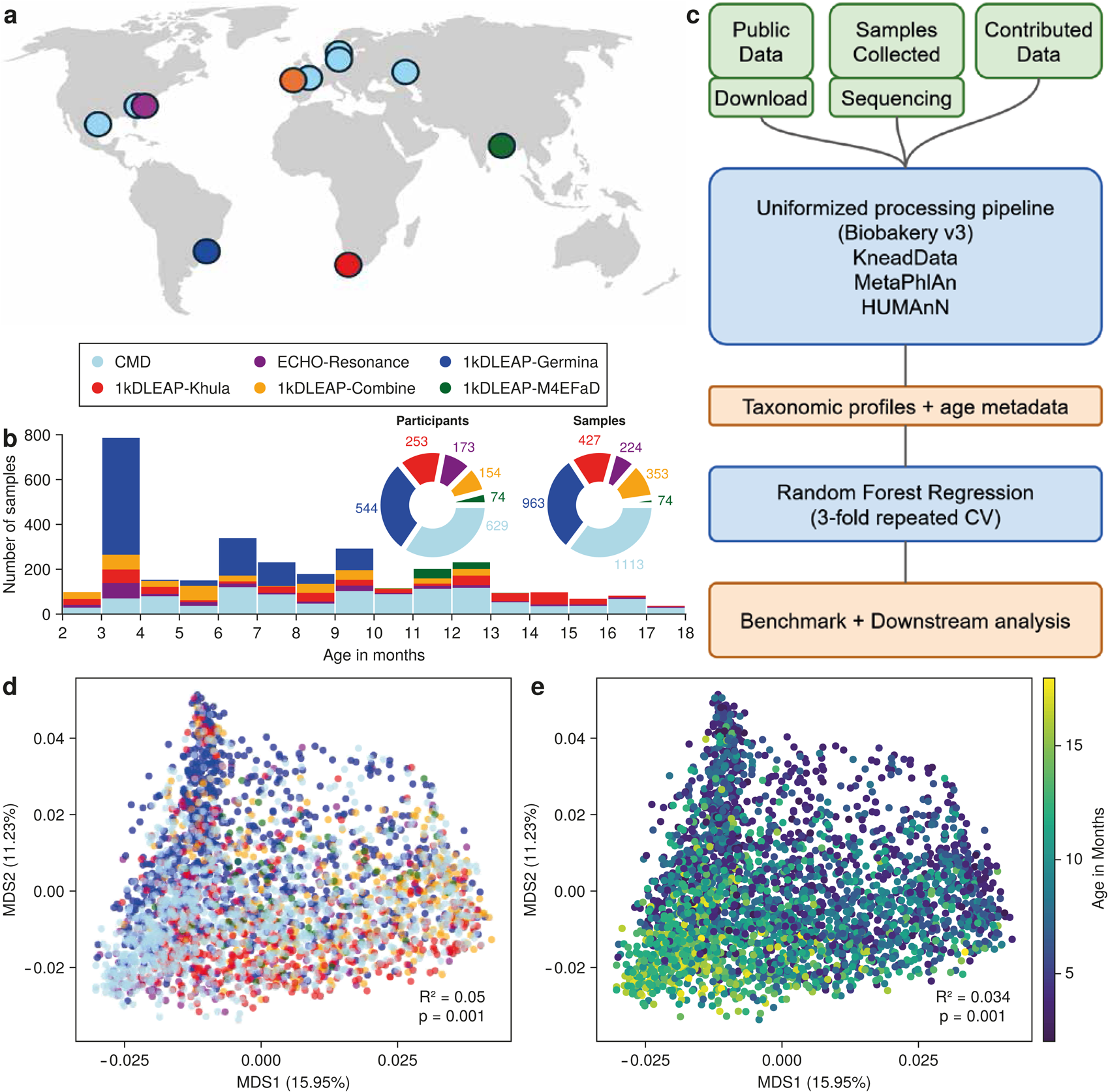
A continuous diversity landscape arises from pooling a large number of globally sampled, uniformly (computationally wise) processed early-life metagenomes. (A) Geographical distribution of sample sources (total n=3,154), color-coded by major data source. (B). Distribution of age at sample collection, binned by months since birth, in the dynamic range of the age model, color-coded by major data source. Donut plot details the total sample contribution by major data source. (C) Overview of methodology, from data acquisition (via sampling, sourcing on public repositories or data collaboration), through the same processing pipeline and downstream statistical analysis. (D-E) NMDS ordination of Bray-Curtis β diversity colored by categorical data source (D) and by continuous age in months (E). Axis percentages denote variance explained by principal coordinates.

**Table 1.**
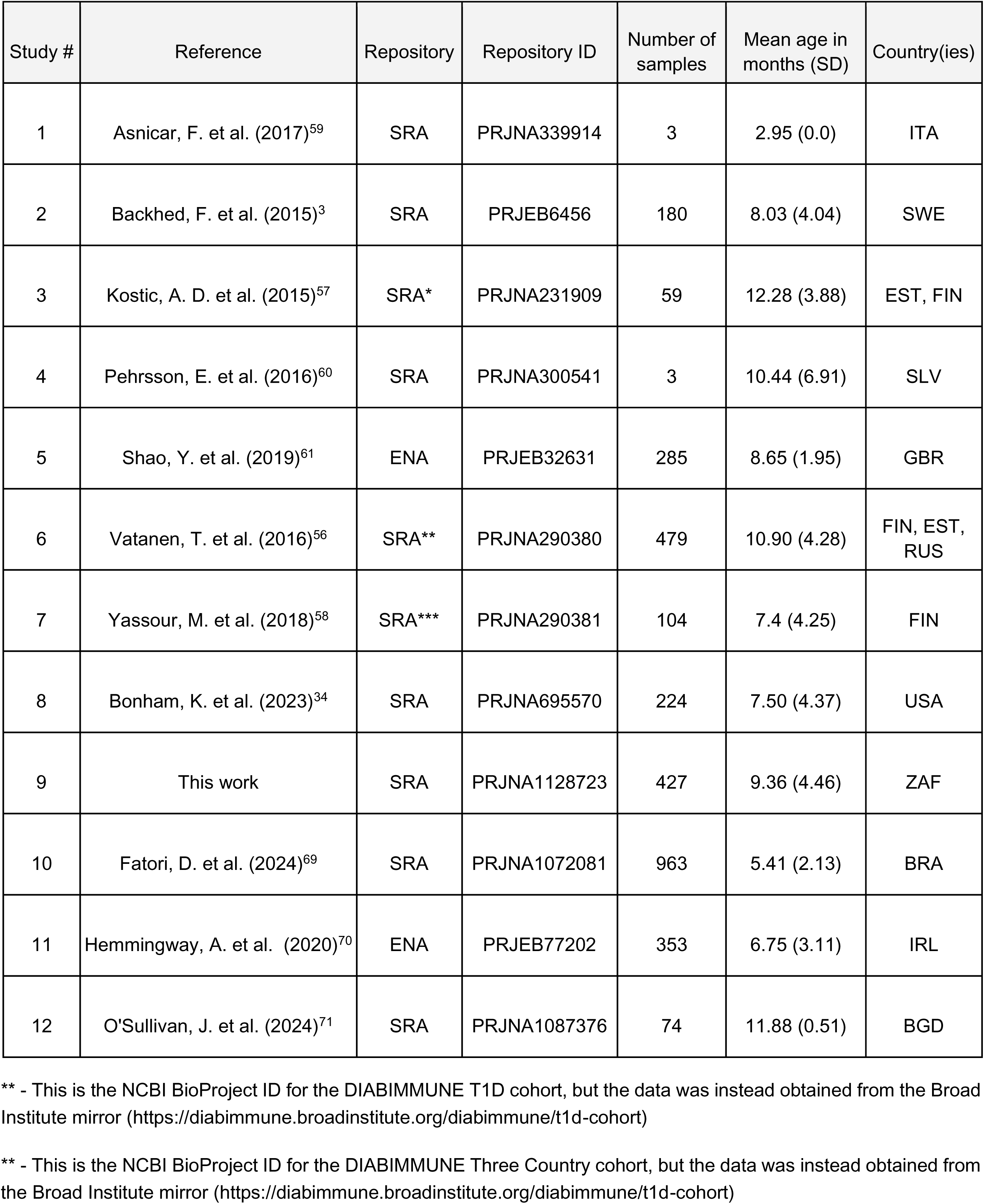
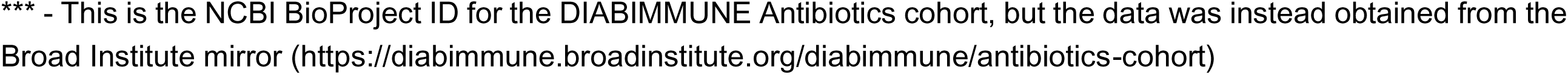
Sources of data for the pooled analysis.

### Harmonized computational processing provides a continuous diversity landscape

After processing all sequence data using the same bioinformatics pipeline (BioBakery V3, **Fig. 1C**), we pooled all community profiles for the downstream analyses. To quantify the variation in gut microbial taxa associated with both age and data source, we used permutational analysis of variance (PERMANOVA) accounting for those factors (**Fig. 1D-E**). Sample group (source) and age explained 5.03% (p = 0.001) and 3.38% (p = 0.001) of the variance, respectively. In a multivariable analysis combining both factors, age still explained 2.28% (p = 0.001) of the variance after accounting for the data source contribution.

### Pooled metagenomes predict age with high resolution

To assess the predictive potential of gut taxonomic profiles for the chronology of gut development, we trained a 5-fold cross-validated (CV) random forest (RF) model on features derived exclusively from the community composition obtained from shotgun metagenomic sequencing. Our inputs were the relative abundances of species present in at least 5% of samples, alongside the α-diversity estimated as the Shannon index. After removing samples with no reads assigned to at least one of the prevalence-filtered species, our analysis comprised 3,153 samples (∼630 per fold) and 149 species. Our model targeted continuous age as a univariate regression output and generated validation-set predictions that reach a root mean square error of cross-validation (RMSECV) of 2.56 months (16.0% of the effective dynamic range, 64.1% of output SD) and a Pearson correlation of 0.803 with the ground truth values, on a 100x repeated 5-fold CV setting (**Fig. 2A**).

**Figure 2.**
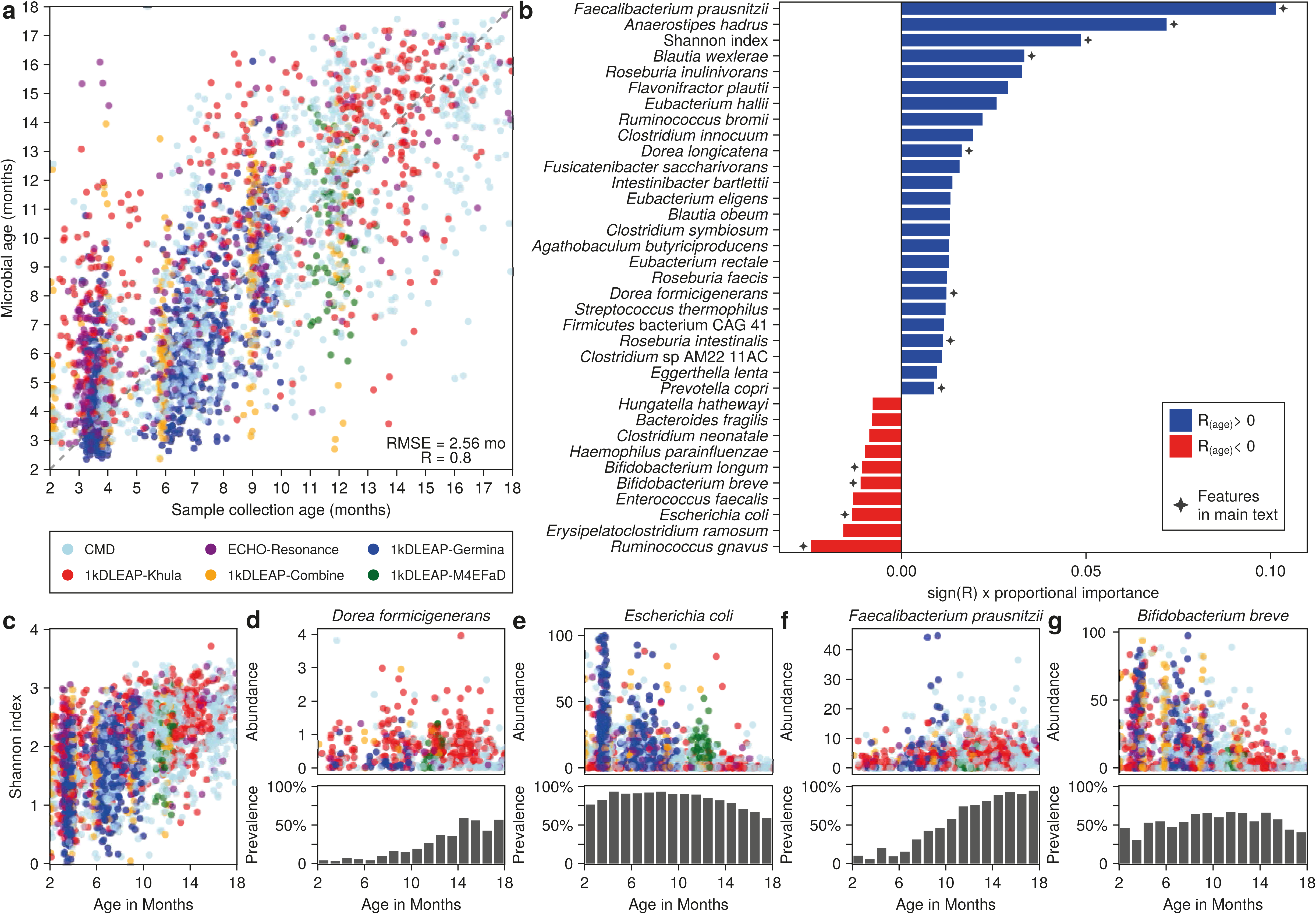
Gut microbial taxon abundances from shotgun metagenomics predict host age with high accuracy in early infancy. (A) Validation-set predicted ages versus ground-truth ages for all samples, color-coded by major data source. (B) Directional importances of top predictive features measured as mean decrease in impurity (MDI) for the trained RF models, multiplied by sign of correlation between predictor and outcome. Absolute values in the x-axis represent a proportion of the total fitness-weighted importance assigned to features. (C) Shannon index with respect to host age, color-coded by major data source. (D-G) Relative abundances color-coded by major data source and average month-by-month prevalences of the indicated important species, *D. formicigenerans* (D)*, E. coli* (E)*, F. prausnitzii* (F), and *B. breve* (G), with respect to host age.

### Changing taxa show feeding transitions and dietary exposures

To derive biological insight from the trained models, we analyzed the fitness-weighted variable importances on the cross-validated models, producing a list of top predictive features (**Fig. 2B**). The 35 highest ranking predictors (23.3% of inputs) were responsible for 70% of the cumulative weighted variable importance. Among those, 25 (71.4%) were positively correlated with age (mean R_(age)_ = 0.18, SD = 0.12), with the remaining 10 (28.6)% negatively correlated to age (mean R_(age)_ = −0.11, SD = 0.07, Supplementary Fig. 2). α-diversity measured as the Shannon index was the third most important predictor (4.86% of total importance, R_(age)_ = +0.52, **Fig. 2C**). All but one of the top predictive taxa (97%) were present in every major cohort (*see* **Methods**), with only *Roseburia intestinalis* remaining undetected in the 1kDLEAP-M4EFaD samples. Additionally, there were several examples of site-biased or site-specific importances. For instance, *Dorea longicatena* and *Dorea formicigenerans* (**Fig. 2D**) were elevated in the South African cohort, and *Escherichia coli* (**Fig. 2E**) was elevated in the Brazilian cohort. Most of the top predictive taxa are species consistently prevalent across all cohorts, indicating that the relevant predictors are robust indicators of age across diverse populations, overcoming population-specific effects.

Across all cohorts, *Faecalibacterium prausnitzii* (**Fig. 2F**) and *Anaerostipes hadrus* were the taxa with the greatest importance scores for age prediction, accounting for 17.3% of the total weighted variable importance, together. Individually, those species positively correlate with age in our dataset (respectively, +0.41 and +0.32). The opposite trend is observed in another key group of predictors that include *Bifidobacterium longum* and *Bifidobacterium breve* (**Fig. 2G**), with 2.2% combined importance, exhibiting negative prior correlations with age (respectively, −0.14 and −0.14). The presence of certain species in the family Lachnospiraceae previously tied to developmental outcomes, such as *Ruminococcus gnavus* and *Blautia wexlerae*^34^ is also noteworthy as a cluster of high-importance predictors of age. The former follows the same trend as the *Bifidobacterium* spp. (2.5% of total importance, R_(age)_ = −0.063, p = 0.001), in agreement with previous studies^35^.

### Learned gut microbial patterns generalize across different sites

To evaluate the generalizability of our model across different data sources and test the predictive ability of each data source toward age, we performed a leave-one-datasource-out cross-validation (LOOCV) experiment. LOOCV yielded an average RMSE of leave-one-out cross-validation of 3.03 +- 0.63 months (Supplementary Table 1, Supplementary Fig. 3). We hypothesized that this generalizability resulted from combined effects from abundance trends and underlying prevalence trends (**Fig. 2D-G**, Supplementary Fig. 2). This would mean that predictors would be important, in part, because they would appear and disappear from the infant gut following similar trends, regardless of geographical origin.

By grouping a subset of our samples by location - Baltic countries, United States and South Africa - and binning them by age (in months), we computed monthly prevalences for the 34 top taxonomic predictors of gut chronology. Strikingly similar patterns of succession emerged between all tested locations (**Fig. 3**), evidenced by whole-matrix mean prevalence correlations: Baltic/USA = 0.799; USA/SA = 0.750; Baltic/SA = 0.749. This consistency suggests that many of the succession patterns identified by our model are likely universal, transcending local environmental influences.

**Figure 3.**
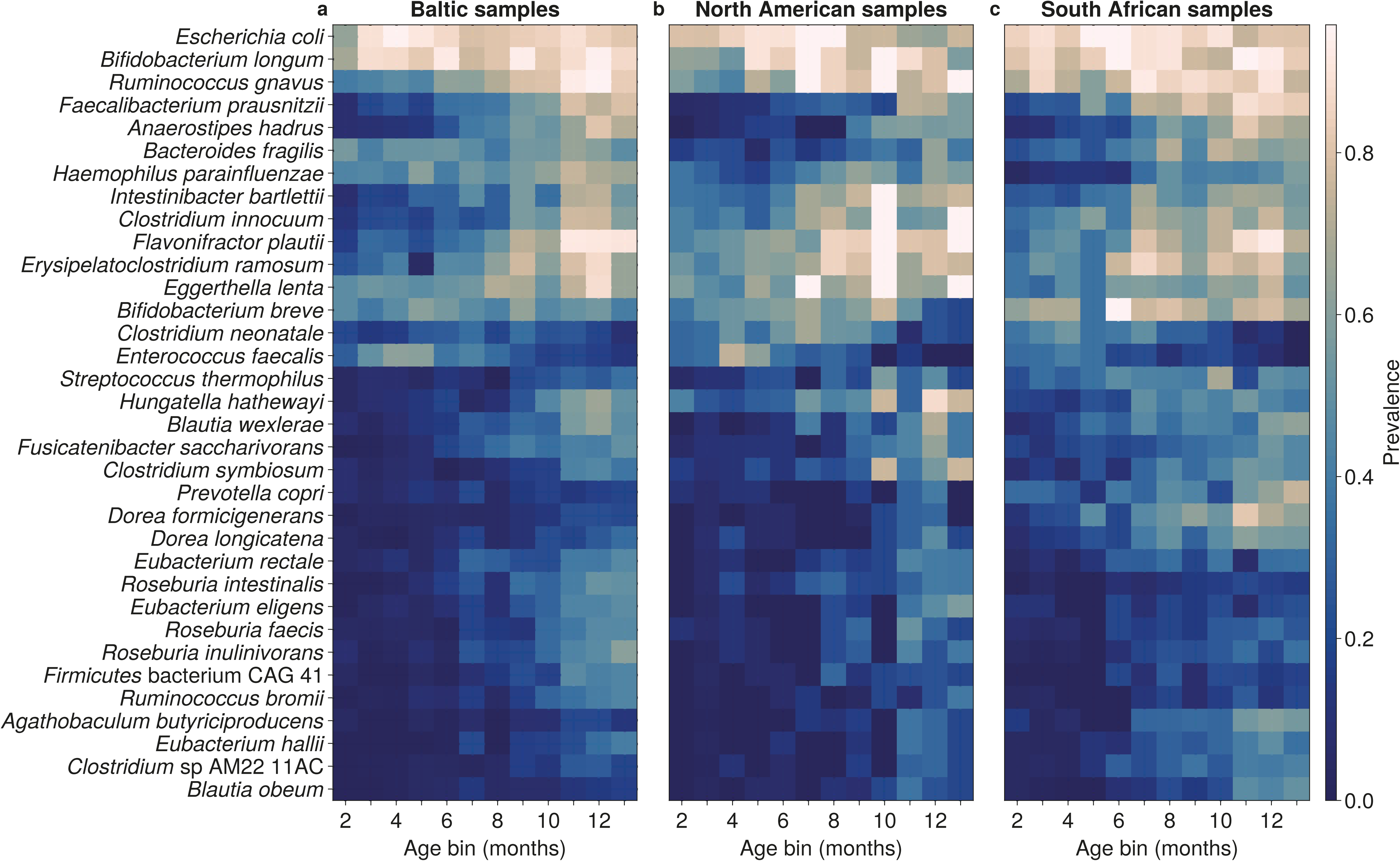
Temporal succession patterns for a common core of age-predictive taxa generalize beyond geographical boundaries. Heatmaps of average taxon prevalence for each of the top 30 predictive species highlighted in Fig. 2. Species are ordered vertically by minimal-distance hierarchical clustering. Samples are binned horizontally from 2 to 13 months. Each cell represents the mean prevalence of that species in the samples collected on that specific month. Panels represent samples belonging to (A) Baltic countries (FIN, EST, RUS, SWE); (B) the United States (USA) and (C) South Africa (ZAF).

Hierarchical cluster analysis of the binned prevalence time series revealed one large universal cluster of species and succession patterns containing 18 (53%) of the top 34 taxonomic predictors, which correlated highly between sites, along with smaller clusters of decentralized patterns. Representatives of the larger, common core are species as mentioned above, such as *F. prausznitzii,* positively correlated with the outcome on all three cases, alongside early colonizers such as *E. coli* (1.3% of total importance, R = −0.25 with age), that follow the opposite pattern consistently on the three sites. Among the divergent cluster, besides the aforementioned *Dorea* genus (*D. longicatena* and *D. formigicerans,* 2.8% combined importance) in South Africa, we identified taxa such as *Prevotella copri* (0.9% of total importance, R = +0.22 with age), which exhibit distinct abundance and prevalence patterns between westernized and non-westernized populations^36^.

### Enzyme changes in the first year corroborate prior studies

We hypothesized that, as was the case with taxonomic composition, the functional composition in terms of microbial metabolic enzymes would change similarly between sites. Utilizing longitudinal samples in the South African cohort, we measured the consistency of the direction of EC abundance transitions between earlier and later samples from the same subject using a Transition Score (TS, *see **Methods***). We then selected the top hits in both directions - later enrichment (highest scores) and later depletion (lowest scores), and stratified their abundances into the corresponding top predictive taxa (**Fig. 4**).

**Figure 4.**
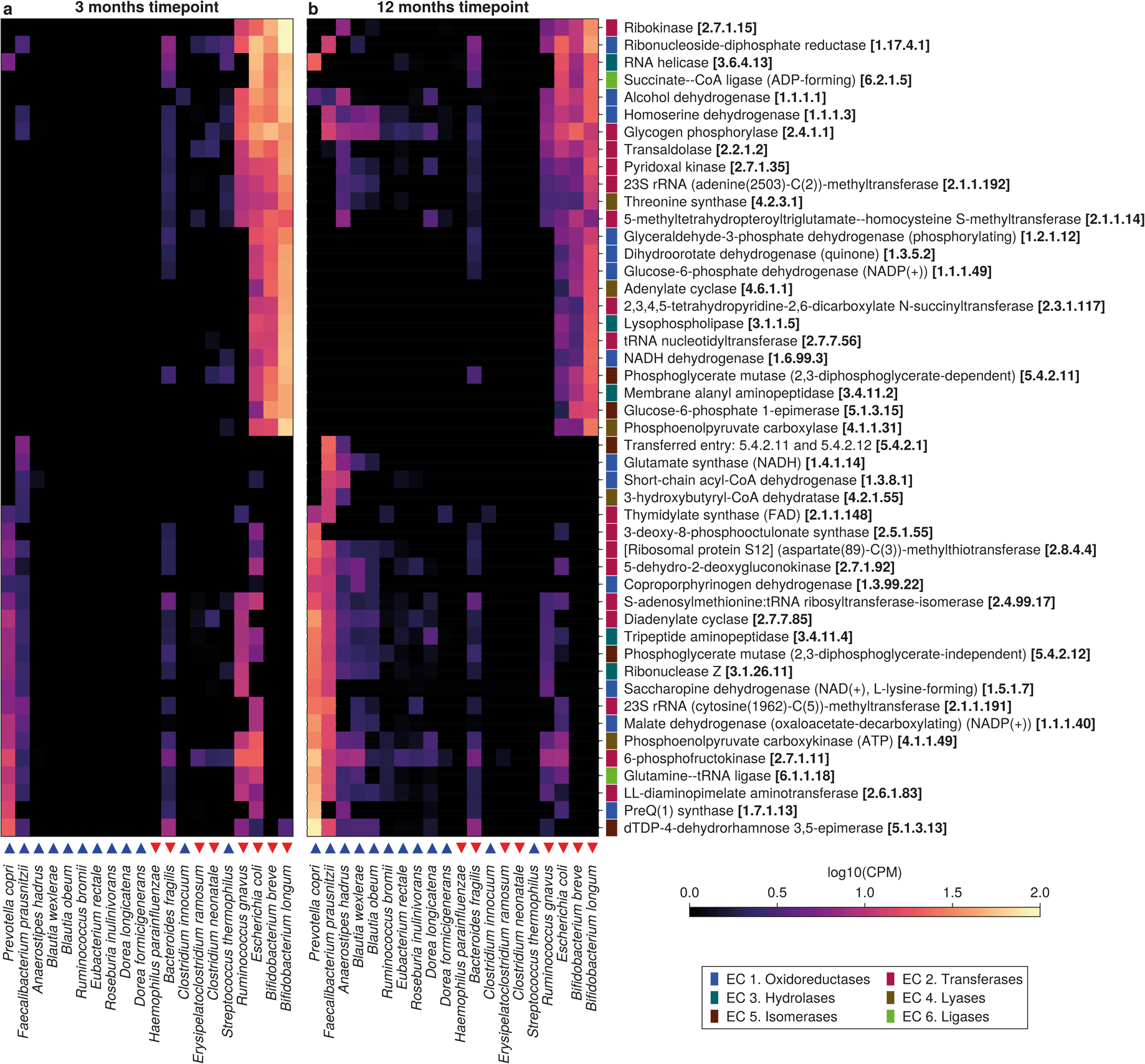
Functional changes are driven by taxonomic changes and centered around diet-associated pathways. Top 24 increasing and top 24 decreasing ECs (in community-wide abundance), stratified in a selected subset of the top taxonomic predictors of age. Cell colors reflect taxon-stratified EC abundance on younger (A) and older (B) samples, measured in log_10_ CPM (counts per million reads). Blue and red triangles indicate species that increase and decrease in abundance in the first year of life, respectively.

The lowest-scored EC (decreasing on most subjects) was transaldolase (2.2.1.2), with a TS of −0.84 and a variation of −86.74 ± 11.46 counts per million reads (CPM). It is followed by succinate-CoA ligase (ADP-forming, 6.2.1.5) and pyridoxal kinase (2.7.1.35), both with a TS of −0.81 and variations of −119.89 ± 20.15 CPM and −67.40 ± 11.44, respectively. The expanded list of stratified ECs decreasing in abundance with age was dominated by functions associated with *B. longum, B. breve, R. gnavus* and *E. coli*, consistent with the aforementioned depletion of those species along the first year of life. That group of species and the highlighted functions account for a consistent average fold change of −0.46 ± 0.01 log_10_ CPM between younger and older samples.

The highest-scored ECs (increasing on most subjects) were [ribosomal protein S12] (aspartate(89)-C(3))-methylthiotransferase (2.8.4.4, TS = +0.84, Δ = +53.89 ± 9.49 CPM), and coproporphyrinogen dehydrogenase (1.3.99.22, TS = +0.79, Δ = +31.54 ± 5.18 CPM). Stratification of the ECs that increase in abundance with age is more diverse, and contains ECs assigned to a wider array of fastidious anaerobes: *F. prausnitzii*, *A. hadrus*, *B. wexlerae*, *Blautia obeum, D. longicatena* and *P. copri*. Combined, highlighted functions assigned to those species exhibit an average fold change of +0.99 ± 0.10 log_10_ CPM between younger and older samples.

When compared to the results published by Vatanen and colleagues^37^, our list of the top 1.5% increasing or decreasing ECs (**Fig. 4**) contains 11 (27.5%) of the previously-reported transitioning ECs. This overlap between the results happened on both major trend clusters, as exemplified by the previously reported decreases in ribokinase (2.7.1.15, TS = −0.73, Δ = − 155.44 ± 22.25 CPM) and transaldolase or the increase in 6-phosphofructokinase (2.7.1.11, TS = +0.59, Δ = +102.66 ± 25.10 CPM). Furthermore, we identified transitioning ECs not previously reported. In this group of novel ECs, notable variations were the decrease in pyridoxal kinase and the increase in malate dehydrogenase (1.1.1.40, TS = 0.66, Δ = +39.62 ± 8.50 CPM).

## Discussion

In this study, we show that the succession of a small number of key taxa in the early-life gut microbiome follows common patterns, even across various geographical and socioeconomic settings. These patterns are strong and consistent enough to be learned by our microbiome age model, allowing it to generalize beyond individual cohort boundaries. One of the main reasons why we were able to build such a robust model was our large-scale pooling strategy, which enabled us to sample diverse backgrounds in, for example, dietary practices and diet composition, an exposure strongly reflected on the learned patterns. As a result, we captured a broad and representative spectrum of microbial profiles, enhancing the robustness of our model towards regional variations, considered a key obstacle to the generalization of microbiome-based models for a variety of phenotypes^38^.

Most studies to date characterized microbiome age using taxonomic classifications from amplicon sequencing of the 16S rRNA gene. Some of the limitations associated with this sequencing technique are the biases introduced by the choice of primers and target region for the experiment, and substantially reduced taxonomic resolution^23–25^. In our work, by building a model using well-defined species identified by metagenomic sequencing, rather than solely relying on 16S rRNA sequencing, we leveraged the ability of the metagenomic approach to sample all genes in a complex sample. The bacterial genes themselves are too highly dimensional and sparse to act as raw simultaneous inputs to multivariable predictive models, but, when processed, allow for the identification of a broader array of taxa at a higher resolution when compared with the depth of information offered by 16S rRNA gene sequencing^23^. Additionally, through the identification of the functional pathways to which those genes belong, we can get a better understanding of how the functional repertoire of the microbial communities evolved with age.

Importance analysis of the fitted random forest models revealed that the main age predictors were the taxa involved in the microbiome’s natural succession influenced by key events such as changes in diet. For example, *F. prausnitzii* and *A. hadrus are* important age predictors in the first two years of life. Those taxa are butyric acid producers^39^ that usually appear with the cessation of breastfeeding, which marks the transition to a Firmicutes-dominated gut characterized with increased production of short-chain fatty acids (SCFA)^40,41^. The same phenomenon explains the learned importance of known metabolizers of human milk oligosaccharides, namely *Bifidobacterium* spp.^42^, characteristic of the early stages of infancy, especially in locations where exclusive breastfeeding is prevalent. Alongside these taxa, the Shannon index (alpha diversity) also emerged as an important predictor. This was expected, as microbial diversity in the gut increases with age in early infancy^25^. Many of the top predictive taxa showed similar succession patterns during the first 13 months of life (**Fig. 3**) across all tested geographical sites (USA, Europe, South Africa), despite significant socioeconomic differences. This suggests that there is a strong, consistent, and machine-learnable pattern for determining age based on microbial succession, regardless of metadata variations, among the geographical sites tested in this work.

Our study corroborates a significant portion of the results from a previous study^37^ that also examined temporal transitions in ECs in early life. This implies that age-determining taxa and their functions are consistent across different microbial communities, even with the diverse lifestyles and ethnic backgrounds of the several cohorts sampled^32^. The ECs that showed most change were primarily involved in central carbohydrate metabolisms, many of which are associated with bifidobacteria. For example, *B. breve* utilizes ribokinase (2.7.1.15) to harvest ribose as a carbon source in the early gut^43^, and several *Bifidobacterium* spp. have transaldolase (2.2.1.2)^44,45^. The presence of glycolytic and pentose-phosphate cycle enzymes supports the idea that diet-related transitions, particularly those tied to the intake of complex carbohydrates, are major drivers of age-determining patterns. In this context, one enzyme of particular interest is pyridoxal kinase (2.7.1.35), which plays a role in the GABA synthesis pathway typical of bifidobacteria^46^. Notably, GABA concentrations in infant stool have been associated with behavioral traits in early infancy^47,48^. Our findings suggest a specific functional link of this association between GABA and *Bifidobacterium* spp. that is also related to age, highlighting a pathway that can be a strong candidate for studying behavioral outcomes in the first year of life.

Despite the strong benchmarks reported by our models, there are several limitations that future studies need to address. For example, one key decision in our model development was to exclude all additional participant and biospecimen metadata, using only participant age and microbial data. This decision was made due to the lack of uniformity in metadata collection and annotation across studies. However, previous studies have shown that metadata such as feeding practices^14^, socioeconomic status^49^, delivery mode and gestational age^50^ can enhance the predictive power in microbial-based models. Notably, in our case, including these covariables would have resulted in a significant loss of samples due to missing metadata, which would have compromised the model’s generalizability and made comparative benchmarks unfeasible. Another area of improvement would be to incorporate season as an external effect to model the time-serial succession patterns, accounting^51^ for different hemispheres. It is also worth mentioning that, even though there are many reference genomes for the early-life gut microorganisms, detailed information on their functions and biochemical characteristics is still biased toward a few well-characterized microorganisms^52^. While we were able to corroborate findings from Vatanen et al. (2018) despite the time gap between the studies, this may partly be due to the limited characterization of the annotated functional space.

Studying developmental changes associated with dynamic processes can be challenging without benchmarks or standards that provide expected ranges of values. Given the high dimensional and highly dynamic nature of microbial composition, simple standards such as those used in anthropometrics (e.g., age-standardized Z-scores for length or weight in infants) are not feasible, and studying microbial associations with child development has been challenging without such an agreed upon normative developmental trajectory. The microbiome-age model provided here, built from a diverse and global population of human children provides a model of development that may be deployed to advance our understanding of the gut microbiome in child growth and flourishing.

## Methods

### Sample collection and processing for the Khula cohort

#### Participants and study design

Infants were recruited from local community clinics in Gugulethu, an informal settlement in Cape Town, South Africa as part of an ongoing longitudinal study (most of the enrollment happened prenatally with 38.82% of infants enrolled shortly after birth^33^). The first language of the majority of residents in this area is Xhosa. Study procedures were offered in English or Xhosa depending on the language preference of the mother. This study was approved by the Health Research Ethics Committees (study number: 666/2021). Informed consent was collected from mothers on behalf of themself and their infants. Demographic information, including maternal place of birth, primary spoken language, maternal age at enrollment, maternal educational attainment, and maternal income, was collected at enrollment (**Table 2**).

**Table 2.**
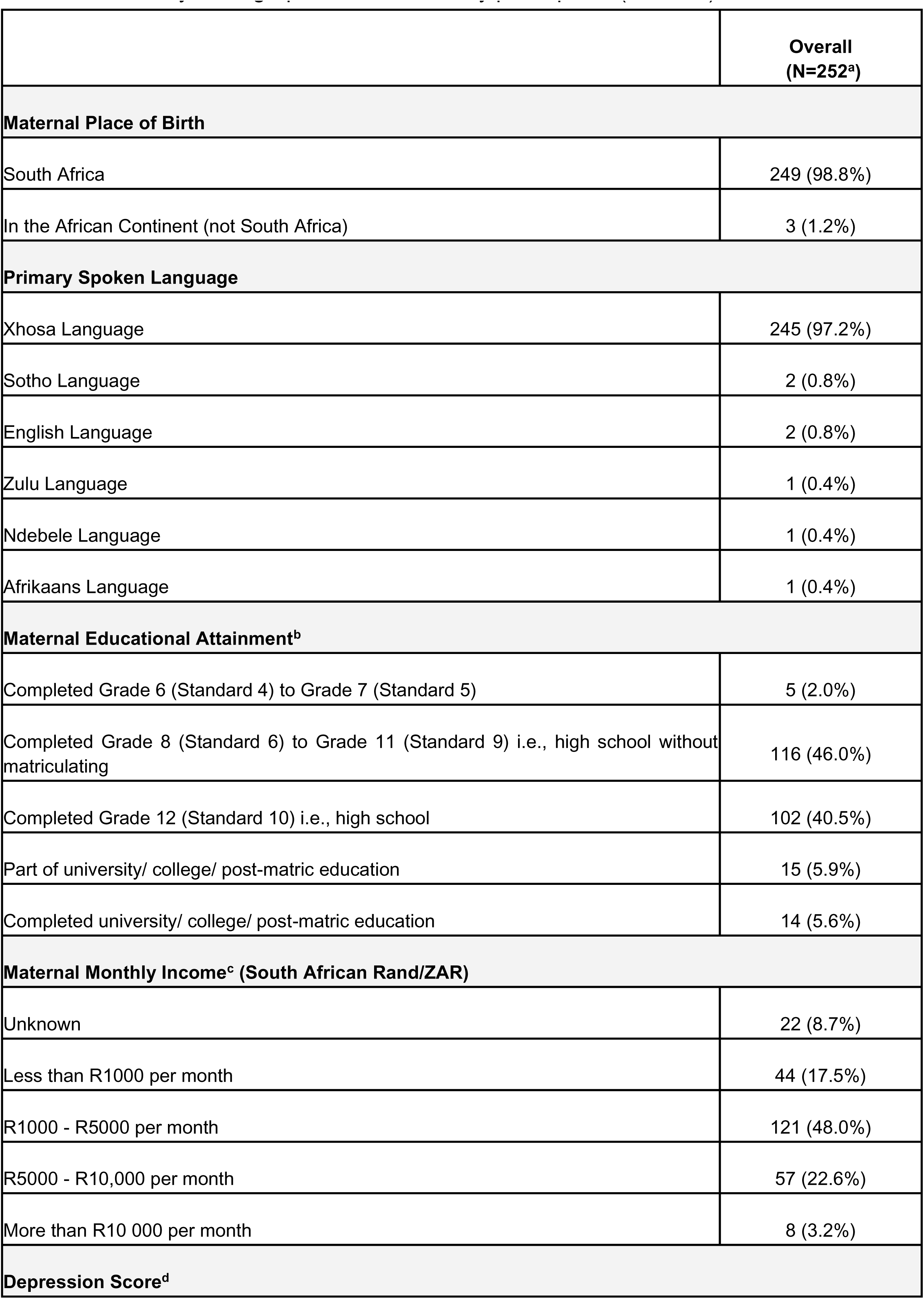
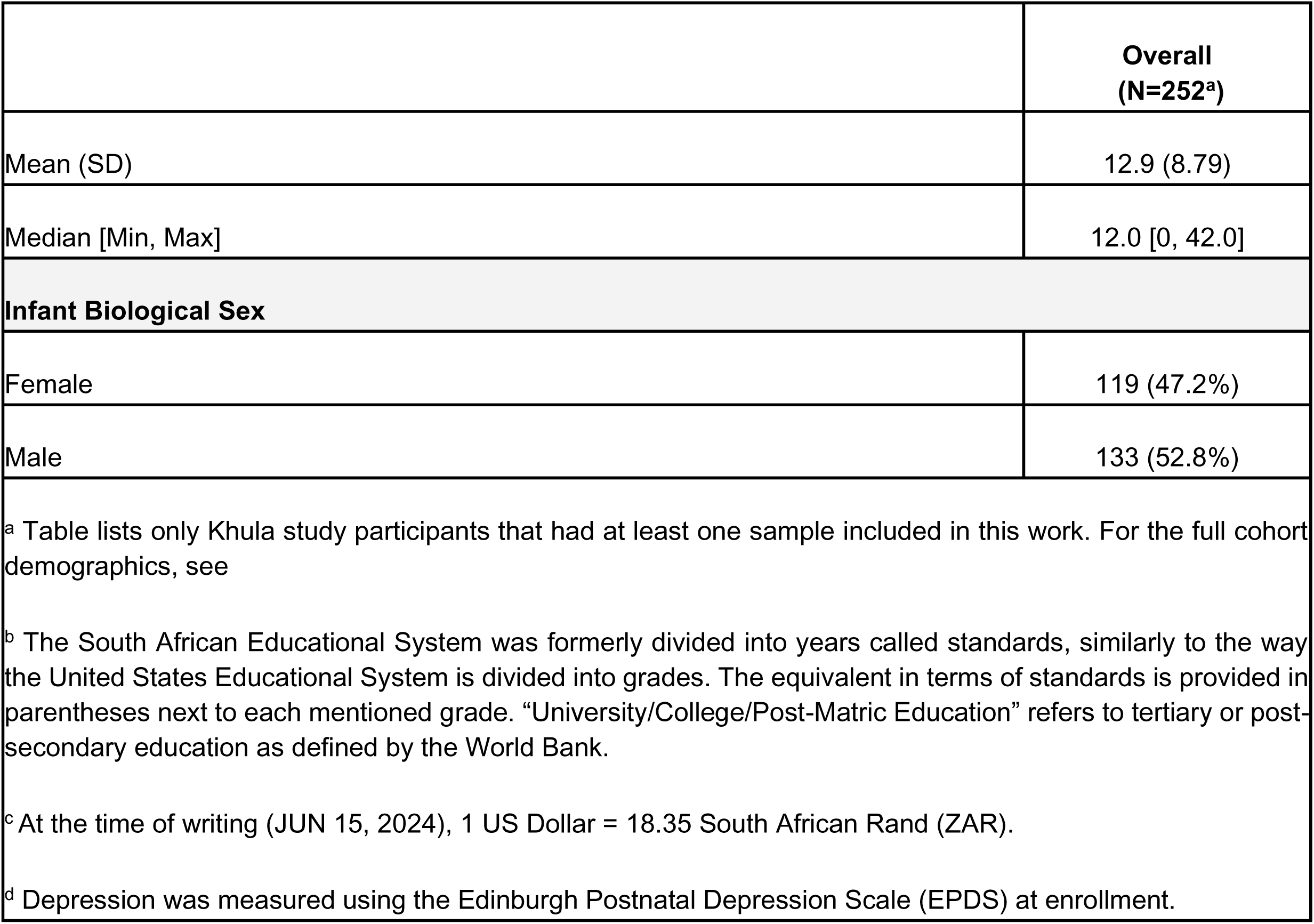
Summary demographics of Khula study participants (mothers)

Families were invited to participate in three in-lab study visits over their infant’s first 18 months of life. At the first in-lab study visit (hereafter Visit 1), which took place when the infants were between approximately 2 and 6 months of age (M=3.63, SD=0.78, range=2.13-5.34), the following data were collected: the infants’ age (in months), sex, and infant stool samples. At the second study visit (hereafter Visit 2), occurring when infants were between approximately 6 months and 12 months of age (age in months: M=8.77, SD=1.39, range=5.38-11.90) and at the third study visit (hereafter Visit 3), occurring when infants were between approximately 12 months and 17 months of age (age in months: M=14.01, SD=1.31, range=11.63-17.97), infant stool samples were collected again. At visits where infants could not donate stool samples on the same day, samples were collected on different days close to the visit date.

#### Sample collection

Stool samples (n=427) were collected in the clinic by the research assistant directly from the diaper and transferred to the Zymo DNA/RNA Shield^TM^ Fecal collection Tube (#R1101, Zymo Research Corp., Irvine, USA) and immediately frozen at −80 °C. Stool samples were not collected if the subject had taken antibiotics within the two weeks prior to sampling.

#### DNA extraction

DNA was extracted at the Medical Microbiology Department, University of Cape Town, South Africa from stool samples collected in DNA/RNA Shield™ Fecal collection tube using the Zymo Research Fecal DNA MiniPrep kit (# D4300, Zymo Research Corp., Irvine, USA) following manufacturer’s protocol. To assess the extraction process’s quality, ZymoBIOMICS® Microbial Community Standards (#D6300 and #D6310, Zymo Research Corp., Irvine, USA) were incorporated and subjected to the identical process as the stool samples. The DNA yield and purity were determined using the NanoDrop® ND −1000 (Nanodrop Technologies Inc. Wilmington, USA).

#### Sequencing

Shotgun metagenomic sequencing was performed on all samples at the Integrated Microbiome Research Resource (IMR, Dalhousie University, NS, Canada). A pooled library (max 96 samples per run) was prepared using the Illumina Nextera Flex Kit for MiSeq and NextSeq from 1 ng of each sample. Samples were then pooled onto a plate and sequenced on the Illumina NextSeq 2000 platform using 150+150 bp paired-end P3 cells, generating on average 24M million raw reads and 3.6 Gb of sequence per sample^53^.

### Public metagenomic data acquisition

Publicly available metagenome metadata was obtained from the CuratedMetagenomicsData database^54^. Database entries considered for inclusion were those annotated as stool samples on the “body_site” property, pertaining to subjects identified as either “newborn” or “child” on the “age_category” property and containing a valid numeric “infant_age” annotation in days. From that set, samples identified as belonging to premature-born children were excluded. We also excluded samples belonging to children suffering from acute infectious conditions - including sepsis - at the time of sample collection. Future T1D-annotated samples, however, (3.9% of the CMD-DIABIMMUNE samples) were not excluded. For the three DIABIMMUNE cohorts, complementary metadata containing harmonized annotation was gathered from the DIABIMMUNE study website and merged with the original set. Sequence data was then downloaded from originally referenced data repositories (**Table 1**).

### Computational processing, analyses and statistics

#### Metagenome processing

For the 1kDLEAP-Khula cohort samples, raw metagenomic sequence reads (Mean = 20.19M, SD = 6.75M reads per sample) were processed using tools from the bioBakery suite, following already-established protocols ^55^. Initially, KneadData v0.10.0 was employed with default settings to trim low-quality reads and eliminate human sequences, using the hg37 reference database.

Subsequently, MetaPhlAn v3.1.0, utilizing the mpa_v31_CHOCOPhlAn_201901 database, was applied with default parameters to map microbial marker genes and generate taxonomic profiles. The taxonomic profiles, along with the same reads obtained in the initial step, were then processed with HUMAnN v3.7 to produce stratified functional profiles. Utilizing this pipeline, the 1kDLEAP-Khula, the ECHO-Resonance^34^ (Mean = 9.34M, SD = 6.75M reads per sample) and the CMD sequence reads (Mean = 15.35M, SD = 13.72M reads per sample) were processed at Wellesley College, USA; the 1kDLEAP-Germina (Mean = 8.32M, SD = 6.48M reads per sample) sequences were processed at the University of Sao Paulo, Brazil; the 1kDLEAP-Combine (Mean = 8.32M, SD = 6.48M reads per sample) sequences were processed at the APC Microbiome Ireland, Ireland; and the 1kDLEAP-M4EFaD (Mean = 41.45M, SD = 6.63M reads per sample) sequences were processed at the Liggins Institute, New Zealand.

#### Sample pooling

Samples were pooled into the same collective dataset and were annotated to differentiate their original data source. For the 4 Wellcome LEAP 1kD studies, every individual study became one separate annotated data source. ECHO-Resonance samples were also annotated as their individual data source. For simplification purposes in downstream analysis, all the CMD-derived samples were annotated as belonging to the same meta-datasource, “CMD.” In analyses that warranted a higher degree of discrimination, we divided this meta-group into two meta-subgroups, “CMD-DIABIMMUNE” (containing 642 samples from Vatanen et al.^56^, Kostic et al.^57^ and Yassour et al.^58^) and “CMD-OTHER” (containing 471 samples from Asnicar et al.^59^, Bäckhed et al.^3^, Pehrsson et al.^60^, Shao et al.^61^).

#### Microbial community analysis

Computational analysis was conducted using the Julia programming language^62^. Microbial community profiles (taxonomic and functional) were parsed and processed using the BiobakeryUtils.jl and Microbiome.jl packages^63^. Principal coordinates analysis (PCoA) with the Bray-Curtis dissimilarity was calculated for all pairs of samples, focusing on species-level classifications, using Distances.jl. Classical multidimensional scaling (MDS) was then performed on the dissimilarity matrix with MultivariateStats.jl. Additionally, permutational analysis of variance (PERMANOVA) was conducted using PERMANOVA.jl.

#### Machine Learning

Machine learning analysis was performed using the MLJ.jl package^64^ and the associated framework. Random forest regression utilized the backend from the DecisionTree.jl package^65^. Linear bias correction was applied to forest outputs when necessary^66^ using GLM.jl^67^. Data visualization was built using the Makie.jl package^68^.

#### Functional Analysis

EC abundance profiles were obtained for each subject of the 1kDLEAP-Khula cohort that had longitudinal samples collected on the 3 month and 12 month timepoints, for a total of 73 sample pairs. Only ECs that could be assigned to at least one detected species were analyzed. ECs were then assigned a transition score (TS) to represent the directionality and consistency of the change in its abundance between the timepoints. For each EC, the TS score was calculated according to the following expression:

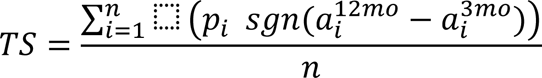

where *n* is the total number of samples; 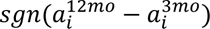 is the sign of the difference in community-wide enzyme abundance for the *i^th^* sample pair between the 12mo and 3mo timepoints; and *p_i_* is a factor that controls for the significance of the EC abundance in either timepoint, according to the expression:

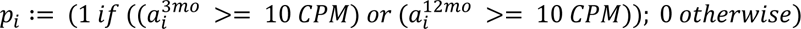

A score close to +1.0 means that the enzyme is consistently increasing from 3 to 12 months, and a score close to −1.0 means that the enzyme is consistently decreasing from 3 to 12 months. After scoring and ranking the ECs, we selected 1.5% of the total scored functions (48 ECs) equally distributed between the highest and lowest-scoring enzymes (24 in each major trend cluster) for stratified functional analysis and visualization.

## Data availability

The processed datasets generated and/or analyzed during the current study have been deposited in Data Dryad under DOI: https://doi.org/10.5061/dryad.dbrv15f9z. The raw sequencing data for the Khula study have been deposited in the NCBI Sequence Read Archive (SRA) under BioProject accession number PRJNA1128723. All other relevant data supporting the key findings of this study and instruction on how to obtain it are available within the article and its Supplementary Information files, or are available from the corresponding author upon reasonable request.

## Code availability

Information for replicating the package environment and code for data analysis and figure generation, as well as scripts for automated download of input files, are available on GitHub at https://github.com/Klepac-Ceraj-Lab/MicrobiomeAgeModel2024 and archived on Zenodo under DOI: https://zenodo.org/doi/10.5281/zenodo.12822332.

## Acknowledgments

We would like to extend our thanks to all the mothers and their infants who generously provided their time and samples for this study. We are also grateful to the dedicated nurses and researchers who recruited participants and collected data, ensuring the success of this project. We would like to express our appreciation to the members of the Huttenhower laboratory for their insightful comments and edits. This research was supported by the Wellcome LEAP 1kD program.

## Author Contributions

Conceptualization - GFB, KSB, VKC; Data curation - GFB, KSB, ID, BCW, DMM; Formal Analysis - GFB, KSB, BCW, SM, FP, NN, PS, DH, AD, RJ, FSM; Funding acquisition - VKC, JMO, KAD, DMM, MEK, ACC, GVP; Investigation - GFB, Methodology - GFB, KSB, CH, FP, SM, VKC; Project administration - VKC; Resources - VKC, KD; Software - GFB, KSB; Supervision - VKC, KSB, FSM, RAB, JMO, CRT, PCBBB, CH, KAD; Validation - GFB; Visualization - GFB; Writing/original draft - GFB; Writing/review & editing - All authors.

## Competing interests

GVP has served as a speaker and/or consultant to Abbott, Ache, Adium, Apsen, EMS, Libbs, Medice, Takeda, developed CME material for Mantecorp and receives authorship royalties from Manole Editors.

## Notes

https://doi.org/10.5061/dryad.dbrv15f9z

https://zenodo.org/doi/10.5281/zenodo.12822332

https://github.com/Klepac-Ceraj-Lab/MicrobiomeAgeModel2024

