## Supplementary Fig. 1 for "Early life microbial succession in the gut follows common patterns in humans across the globe"

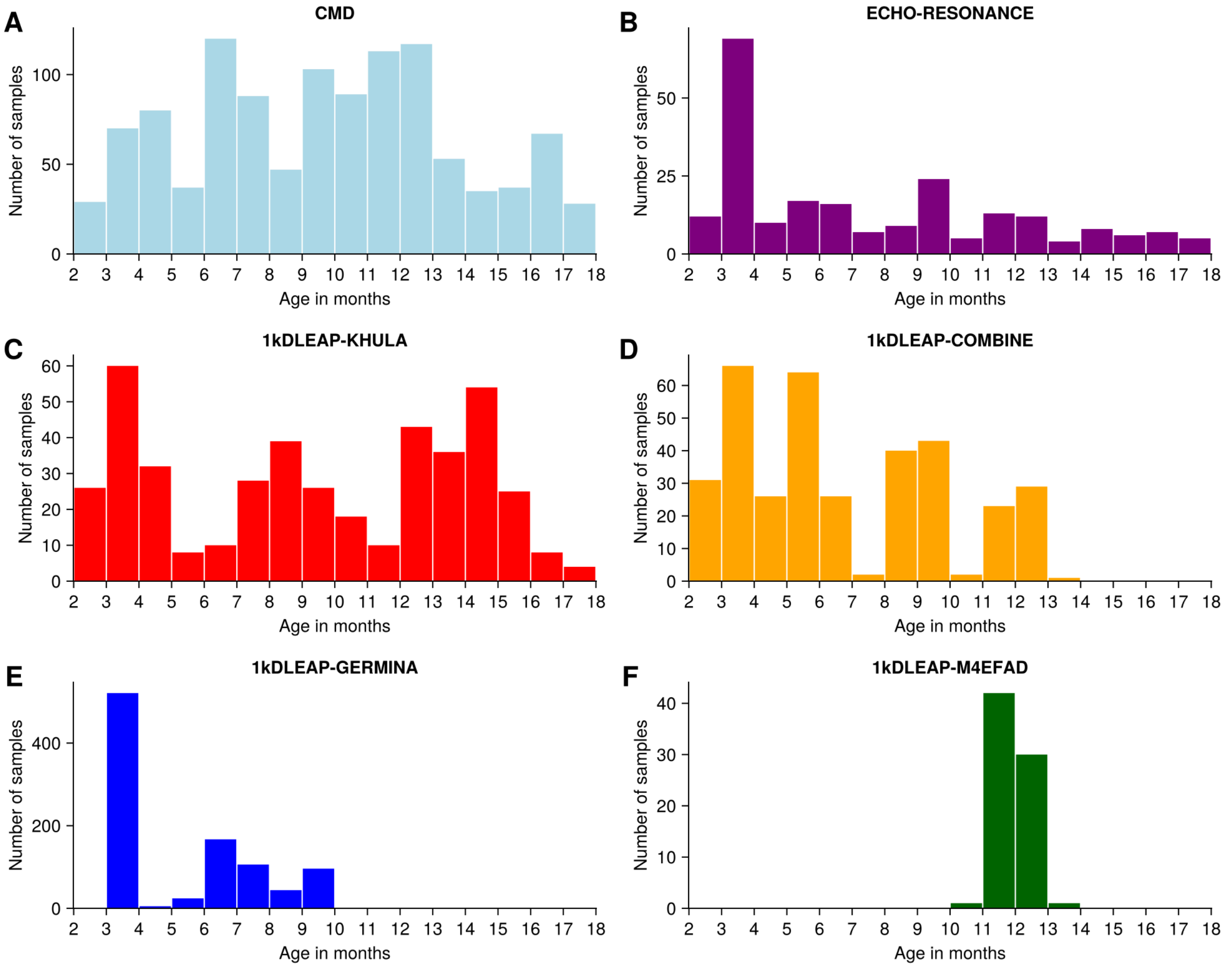


**Supplementary Figure 1. Individual data source contributions to the distribution of ages in the dynamic range.** Individual components of **Main Figure 1B** from each major data source: (A) CMD, (B) ECHO-Resonance, (C) 1kDLEAP-Khula, (D) 1kDLEAP-Combine, (E) 1kDLEAP-Germina, and (F) 1kDLEAP-M4EFaD.
