## Supplementary Fig. 2 for "Early life microbial succession in the gut follows common patterns in humans across the globe"

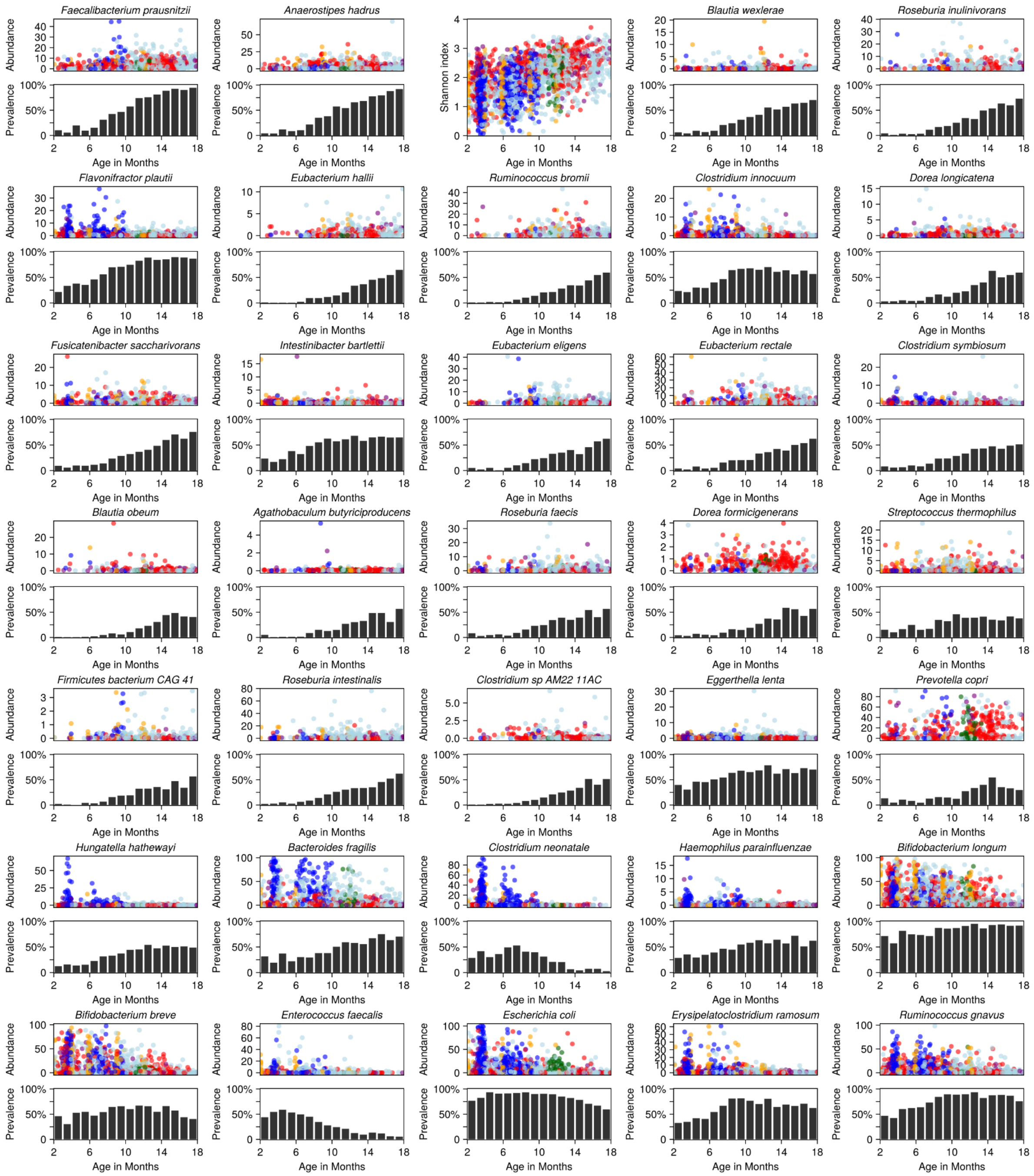


**Supplementary Figure 2. Individual feature association patterns for all 35 top predictive features.** Plots include the α-diversity measured as the Shannon index and the 34 most important species, which combined account for 70% of the cumulative proportional fitness-weighted importance. For each species, plots include abundance (top) and prevalence (bottom) patterns.
