## Supplementary Fig. 3 for "Early life microbial succession in the gut follows common patterns in humans across the globe"

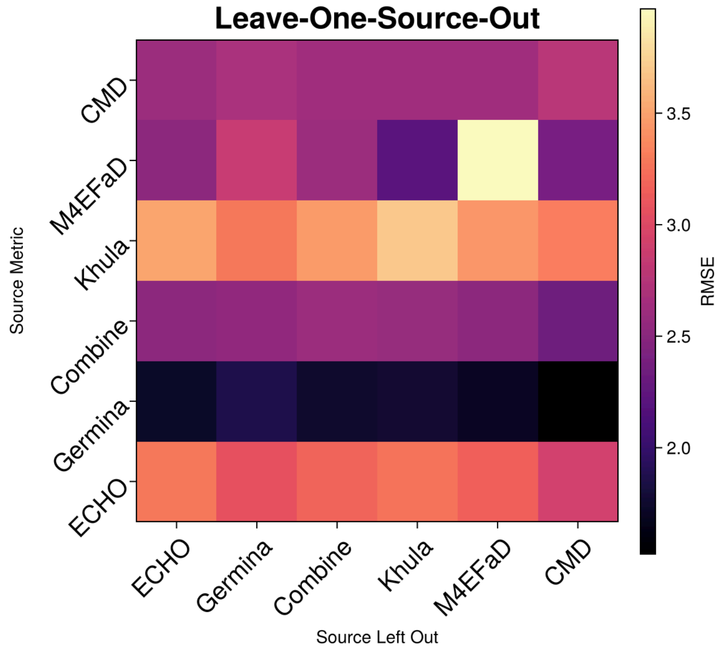


**Supplementary Figure 3. Results matrix for leave-one-out cross-validation experiment.** The Y axis represents the data source for which the RMSE is being measured, and the X axis represents the data source that is being left out as an external test set, versus data sources that participated in internal cross validation.
