## Supplementary Table 1 for "Early life microbial succession in the gut follows common patterns in humans across the globe"

**Supplementary Table 1.** Summary results for leave-one-out cross-validation experiment.

| Data Source left out | Included data sources' validation folds RMSE | left-out data source test RMSE |
|---|---|---|
| CMD | 2.355 ± 0.004 | 2.769 ± 0.002 |
| ECHO-Resonance | 2.510 ± 0.005 | 3.284 ± 0.003 |
| 1kDLEAP-Brainrise | 2.846 ± 0.005 | 1.845 ± 0.002 |
| 1kDLEAP-Khula | 2.420 ± 0.005 | 3.700 ± 0.003 |
| 1kDLEAP-COMBINE | 2.584 ± 0.005 | 2.595 ± 0.002 |
| 1kDLEAP-M4EFaD | 2.5494 ± 0.005 | 3.967 ± 0.005 |
